# Structural and functional characterization of chloroplast ribulose-5-phosphate-3-epimerase from the model green microalga *Chlamydomonas reinhardtii*

**DOI:** 10.1101/2022.09.29.510120

**Authors:** Maria Meloni, Silvia Fanti, Daniele Tedesco, Libero Gurrieri, Paolo Trost, Simona Fermani, Stéphane D. Lemaire, Mirko Zaffagnini, Julien Henri

## Abstract

Photosynthetic carbon fixation relies on Rubisco and ten additional enzymes in the conserved Calvin-Benson-Bassham (CBB) cycle. Epimerization of xylulose-5-phosphate (X5P) into ribulose-5-phosphate (Ru5P) contributes to the regeneration of ribulose-1,5-bisphosphate, the substrate of Rubisco activity. Ribulose-5-phosphate-3-epimerase (RPE) catalyzes the formation of Ru5P but it can also operate in the pentose phosphate pathway (PPP) by catalyzing the reverse reaction. Here, we describe the catalytic and structural properties of the recombinant form of photosynthetic RPE isoform 1 from *Chlamydomonas reinhardtii* (CrRPE1). The enzyme shows catalytic parameters that are variably comparable to those of the paralogues involved in the PPP and CBB cycle but with some notable exceptions. CrRPE1 is a homo-hexamer that exposes a catalytic pocket on the top of an *α*_8_*β*_8_ triose isomerase-type (TIM-) barrel as observed in structurally solved RPE isoforms from both plant and non-plant sources. Despite being identified as a putative target of thiol-based redox modifications, CrRPE1 activity is not altered by redox treatments, indicating that the enzyme does not bear redox sensitive thiol groups and is not regulated by thiol-switching mechanisms. We mapped phosphorylation sites on the crystal structure and the specific location at the entrance of the catalytic cleft supports a phosphorylation-based regulatory mechanism. Overall, this work provides a detailed description of the catalytic and regulatory properties of CrRPE along with structural data, which allow for a deeper understanding of the functioning of this enzyme of the CBB cycle and in setting the basis for possible strategies to improve the photosynthetic metabolism.

## Introduction

The Calvin-Benson-Bassham cycle (CBBC) is the metabolic phase of photosynthesis that allows the endergonic conversion of inorganic carbon (*i.e*., carbon dioxide) into carbohydrates (Emerson and Arnold 1932, Johnson 2016). Fixation of carbon dioxide occurs through the carboxylation of the acceptor ribulose-1,5-diphosphate (RuBP) by the enzyme ribulose-1,5-diphosphate carboxylase/oxygenase (Rubisco) (Benson, Bassham et al. 1952). Subsequently, enzyme-driven regeneration of RuBP is crucial to sustain Rubisco catalysis and requires epimerization of xylulose-5-phosphate (X5P) into ribulose-5-phosphate (Ru5P) at its chiral carbon 3. Epimerization is catalyzed by ribulose-5-phosphate 3-epimerase (RPE, enzyme class 5.1.3.1), which is generally considered to be a metalloenzyme belonging to the ribulose phosphate binding superfamily (Chen, Hartman et al. 1998). In parallel with RPE, the formation of Ru5P is also guaranteed by the isomerization of ribose-5-phosphate (R5P) catalyzed by ribose-5-phosphate isomerase (RPI). Following epimerase/isomerase activities, the regeneration of RuBP is ensured by the activity of phosphoribulokinase (PRK) which catalyzes the ATP-dependent phosphorylation of Ru5P (Jakoby, Brummond et al. 1956).

*Chlamydomonas reinhardtii*, a model unicellular green alga, encodes two RPEs at nuclear genome loci Cre12.g511900.t1.2 (RPE1) and Cre02.g116450.t1.2 (RPE2) (Merchant, Prochnik et al. 2007). RPE1 and RPE2 proteins share 41% sequence identity but RPE2 (Uniprot A8I2N1) is predicted to be cytosolic while RPE1 (Uniprot A8IKW6) is predicted to be localized in the chloroplast. Like in plants, the PPP is duplicated in the chloroplast and the cytosol of Chlamydomonas (Johnson and Alric 2013). This suggests that RPE2 contributes to cytosolic PPP while RPE1 is active in both the chloroplast PPP and the CBBC.

Recent proteomic studies employed Chlamydomonas as a model organism to identify protein targets undergoing post-translational modifications such as thiol-based redox modifications and serine-threonine phosphorylations. The chloroplast RPE1 isoform was identified as a putative target by redox proteomics probing S-glutathionylation and S-nitrosylation sites (Zaffagnini, Bedhomme et al. 2012, Morisse, Zaffagnini et al. 2014), and thioredoxin-mediated disulfide/dithiol exchange (Pérez-Pérez, Mauriès et al. 2017). Likewise, CrRPE1 was found to undergo multiple phosphorylations on three sites at residues Ser50, Thr220, and Ser239 (Wang, Gau et al. 2014). Nevertheless, the effective modulation of protein activity through post-translational modifications is still to be proven.

Over the last decades, several studies characterized RPE isoforms from different organisms including protozoan parasites, photosynthetic and non-photosynthetic bacteria, yeast, mammals, and land plants. Surprisingly, the various RPE showed rather different structural and functional features likely related to an intrinsic dissimilarity reflecting their specific physiological context. At the biochemical level, the sole enzymatic activity monitored so far is the conversion of Ru5P to X5P (*i.e*., the PPP-type activity) and the corresponding Michaelis-Menten constants (*K*_M_) for Ru5P determined for plant and non-plant RPEs revealed large variations, from high to low affinities (0.2-15 mM range) (Kiely, Stuart et al. 1973, Akana, Fedorov et al. 2006). Similarly, turnover numbers (*k*_cat_) are highly variable ranging from ~0.1 to 10^4^ sec^−1^ (Teige, Melzer et al. 1998, Chen, Larimer et al. 1999). Focusing on plant enzymes, the recombinant form of spinach RPE exhibited a *k*_cat_ of 7,100 sec^−1^ and a *K*_M_ for R5P of 0.22 mM (Chen, Hartman et al. 1998, Chen, Larimer et al. 1999), whereas the catalytic parameters of RPE purified from spinach leaf chloroplasts were estimated at 0.138 sec^−1^ and 0.25 mM for *k*_cat_ and *K*_M_, respectively (Teige, Melzer et al. 1998). These values reflected a similar affinity for the substrate while turnover numbers surprisingly deviate by four orders of magnitude. Consequently, derived catalytic efficiencies (*k*_cat_/*K*_M_) strikingly differ from 3.23 × 10^7^ M^−1^ sec^−1^ for the recombinant enzyme to 5.53 × 10^2^ M^−1^ sec^−1^ for the native enzyme. To date, structures of 23 RPE orthologs were experimentally determined and reported in the Protein Data Bank, among which four from *Homo sapiens* (Liang, Ouyang et al. 2011), one from *Streptococcus pyogenes* (Akana, Fedorov et al. 2006), one from *Plasmodium falciparum* (Caruthers, Bosch et al. 2006), and one from *Trypanosoma cruzi* (Gonzalez, Valsecchi et al. 2017). Three RPE structures were determined from photosynthetic organisms: one from the cyanobacterium *Synechocystis sp*. PCC 6803 (Wise, Akana et al. 2004), one cytosolic isoform from *Oryza sativa* (Jelakovic, Kopriva et al. 2003), and one chloroplast isoform from *Solanum tuberosum* (Kopp, Kopriva et al. 1999). In general, RPEs from both plant and non-plant sources fold as an α_8_β_8_ barrel of the triose-phosphate isomerase (TIM-barrel) superfamily. The carboxy-end of the eight parallel β-strands in the barrel defines a shallow surface onto which a metal ion is chelated by a conserved tetrad of two aspartates and two histidines. X5P and Ru5P substrate ligands are typically accommodated in the vicinity of the metal site where reversible epimerization is catalyzed. The identity of the metal was determined for RPE from *Escherichia coli* (Sobota and Imlay 2011) and attributed to iron II (Fe^2+^), while other structural data locate a zinc ion at this position (Jelakovic, Kopriva et al. 2003, Wise, Akana et al. 2004, Akana, Fedorov et al. 2006). Surprisingly, the structural analysis of chloroplast RPE from *Solanum tuberosum* revealed the absence of a metal ion, but instead, a molecule of water was accommodated in the active site center (Kopp, Kopriva et al. 1999).

In the current study, we present the crystal structure of recombinant RPE1 from Chlamydomonas (CrRPE1), determined at a resolution of 1.9 Å. Our model confirms the general features of the epimerase fold and catalytic site. The quaternary structure was examined by means of size-exclusion chromatography and SAXS analysis indicating that CrRPE1 has a hexameric fold. An identical homo-hexameric state was found by immunoblot analysis of algal protein extracts resolved by size-exclusion chromatography. To determine the biochemical properties of CrRPE1, we set up and optimized *in vitro* coupled-enzyme assays to monitor the enzymatic conversion of Ru5P to X5P (*i.e*., PPP-related activity), and for the first time, the epimerization of X5P to Ru5P (*i.e*., CBBC-related activity). Relying on enzymatic assays, we established the kinetic properties of CrRPE1, which has better affinity toward X5P in spite of a more efficient catalytic proficiency with Ru5P as a substrate. A biochemical comparison with the recombinant form of the spinach homologue was also conducted, showing that the two enzymes have similar kinetic features, in sharp contrast to previous studies. Finally, we monitored the sensitivity of CrRPE to both reducing and oxidizing agents and evaluated possible regulatory mechanisms based on redox and phospho-based modifications by mapping putative sites on the molecular structure.

## Material and methods

### Cloning and protein preparation

*Chlamydomonas reinhardtii* open reading frame of gene Cre02.g116450 of the UniProt entry A8IKW6 was searched for chloroplast transit peptide with ChloroP (Emanuelsson, Nielsen et al. 1999), PredAlgo (Tardif, Atteia et al. 2012) and multiple sequence alignments (Figure S1). Nucleotide sequence encoding residues 28 to 265 of the predicted mature protein was PCR-amplified from template AV634644 (HC036a02) of Chlamydomonas expressed sequence tag database (Kazusa). PCR product was digested by *Nco*I and *BamH*I and ligated into pET-3d vector in fusion with an in frame amino-terminal hexa-histidine tag, yielding plasmid pET3d-His_6_-CrRPE1. The plasmid was used to transform *Escherichia coli* BL21 Rosetta2 (DE3) expression strain (Merck, Darmstadt, Germany), grown to exponential phase in 1 L lysogeny broth (LB) medium supplemented with 100 μg mL^−1^ ampicillin. Expression was induced by addition of 0.2 mM isopropyl-β-D-thiogalactopyranoside (IPTG) for 16 h at 30 °C. Cells were harvested by centrifugation, resuspended in 30 mM Tris-HCl, pH 7.9 (buffer A) and lysed by sonication. Clarified lysate was loaded on 3 mL Ni-NTA resin, washed with buffer A supplemented with 30 mM imidazole and step-eluted with buffer A supplemented with a final imidazole concentration of 250 mM. Eluate was desalted on PD-10 column preequilibrated with 30 mM Tris-HCl, pH 7.9 and concentrated by ultrafiltration to 5-10 mg mL^−1^. The molecular mass and purity of recombinant protein were examined by SDS-PAGE and the resulting homogeneous protein solutions were stored at −20 °C. Protein identity was assessed by MALDI-TOF mass spectrometry. Protein concentration was determined spectrophotometrically using a molar extinction coefficient at 280 nm of 14,105 mM^−1^ cm^−1^ and a molar mass of 26,772.9 g mol^−1^.

### Analytical size-exclusion chromatography

Recombinant CrRPE1 was injected on a Superose 6 Increase 10/300 GL column (GE Healthcare, Chicago, Illinois) and isocratically eluted in 50 mM Tris-HCl (pH 7.5) and 150 mM KCl at a 0.5 mL min^−1^ flow rate. The column was calibrated with standard globular proteins (Bio-Rad, Hercules, USA), namely bovine thyroglobulin (670 kDa), bovine γ-globulin (158 kDa), chicken ovalbumin (44 kDa), horse myoglobin (17 kDa), and vitamin B12 (1.35 kDa).

### *In vitro* reporter assays of CrRPE1 activity

The epimerization of Ru5P into X5P (*i.e*., PPP-related activity) was measured using a 4-step coupled assay as previously described (Nowitzki, Wyrich et al. 1995) (Figure S2), with minor modifications. As reporter enzymes we used the recombinant transketolase and triose-phosphate isomerase from Chlamydomonas (CrTK and CrTPI, respectively) (Zaffagnini, Michelet et al. 2014, Pasquini, Fermani et al. 2017), and the rabbit muscle α-glycerophosphate dehydrogenase (α-GDH; Sigma-Aldrich, Saint-Louis, USA). Prior to the activity assays, CrRPE1 was separated by size-exclusion chromatography and eluted fractions corresponding to the hexameric form were pooled and desalted in 30 mM Tris-HCl (pH 7.9). The catalytic activity was assayed spectrophotometrically at 25 °C in a reaction mixture containing 50 mM Tris-HCl (pH 7.9), 15 mM MgCl_2_, 0.1 mM thiamine pyrophosphate (TPP), 0.1% bovine serum albumin, 1 μM CrTK, 4 nM CrTPI, 2 units mL^−1^α-GDH, 2 mM ribose-5-phosphate (R5P), 0.25-4 mM Ru5P, and 0.2 mM reduced nicotinamide adenine dinucleotide (NADH). The reaction was initiated by the addition of CrRPE1 and the enzymatic activity was measured by following the decrease in the absorption at 340 nm using a Cary60 UV/Vis spectrophotometer (Agilent Technologies). Previous studies highlighted interferences in the assay derived from traces of X5P in the Ru5P powder and from traces of epimerase activity in the auxiliary enzymes from commercial sources (Wood 1979). To establish whether NADH oxidation was effectively dependent upon CrRPE1 catalysis (*i.e*. X5P formation and subsequent reactions catalyzed by CrTK, CrTPI and α-GDH; Figure S2), we measured the consumption of NADH in the absence of CrRPE1 and found a limited NADH oxidation corresponding to 5% of the initial NADH (~10 μM) that we ascribed to X5P contamination in the Ru5P solution.

To measure the CBBC-related activity, namely the conversion of X5P to Ru5P, we employed a newly developed 4-step coupled assay involving three reporter enzymes: the recombinant form of phosphoribulokinase from Chlamydomonas (CrPRK, (Gurrieri, Del Giudice et al. 2019)), and commercial pyruvate kinase (PK) and lactate dehydrogenase (LDH) from *Saccharomyces cerevisae* (Sigma-Aldrich, Saint-Louis, USA) (Figure S2). The catalytic activity was measured at 25 °C in a reaction mixture containing 50 mM Tris-HCl (pH 7.9), 10 mM MgCl_2_, 2 mM adenosine triphosphate (ATP), 2.5 mM phosphoenolpyruvate (PEP), 5 units mL^−1^ PK, 6 units mL^−1^ LDH, 6 units mL^−1^ CrPRK, 0.075-1.5 mM X5P, and 0.2 mM reduced nicotinamide adenine dinucleotide (NADH). The reaction was initiated by the addition of CrRPE1 and monitored at 340 nm using a Cary60 UV/Vis spectrophotometer (Agilent Technologies). CrRPE1-dependent enzymatic activities were determined after subtracting background activities measured in the absence of CrRPE1.

### Protein stability and redox sensitivity

The stability of CrRPE1 was assessed by incubating the enzyme (5 μM) at 4 °C or 25 °C in 30 mM Tris-HCl (pH 7.9). At the indicated times, aliquots (1–2 μl) were withdrawn to carry out activity measurements as described above. The redox sensitivity of CrRPE1 (5 μM) was evaluated by incubating the enzyme in the presence of reducing agents (10 mM 2-mercaptoethanol or 10 mM DTT) or oxidizing molecules such as nitrosoglutathione (GSNO), 2 mM oxidized glutathione (GSSG), or 2 mM hydrogen peroxide (H_2_O_2_), or 10 mM oxidized DTT in the absence/presence of recombinant chloroplast TRX f2, m, x, y, and z from Chlamydomonas, (5 μM each) (Lemaire, Tedesco et al. 2018, Marchand, Fermani et al. 2019). All incubations were carried out at 25 °C in 30 mM Tris-HCl (pH 7.9). Control experiments were performed by incubating the enzyme in the presence of buffer alone. At the indicated times, aliquots (1–2 μl) were withdrawn to carry out activity measurements as described above.

### Crystallization, structure determination and model refinement

100 nL of pure CrRPE1 concentrated at 9.5 mg mL^−1^ were mixed with 100 nL of each of the 384 precipitant conditions of the joint center for structural genomics sparse-matrix screen for crystallization (Qiagen, Hilden, Germany) (Lesley and Wilson 2005). Condition 53 of screen III (160 mM Calcium acetate; 80 mM Sodium cacodylate pH 6.5; 20% glycerol; 14.4% polyethylene glycol 8,000) yielded macled rods of 100 μm length. Crystals were flash-frozen in cryo-loops and tested for diffraction at ESRF beamline ID30A-3 (Grenoble, France). 7,200 frames of 0.05° tilt each were collected for a complete dataset of space group P2_1_ (Table 1). Indexation, integration, and scaling were performed with XDS (Kabsch 2010) on the beamline software EDNA (Incardona, Bourenkov et al. 2009). Phases were determined by molecular replacement with PHASER-MR (McCoy, Grosse-Kunstleve et al. 2007) from protomers of CrRPE1 as modelled by homology with PHYRE2 prediction algorithm (Kelley, Mezulis et al. 2015). Twelve subunits were searched for, to respect a solvent content of 48.8 % in the asymmetric unit. Initial density map and structural model were interpreted by iterative cycles of automated building with AUTOBUILD (Terwilliger, Grosse-Kunstleve et al. 2008), manual building with COOT (Emsley, Lohkamp et al. 2010) and refinement with PHENIX.REFINE (Afonine, Grosse-Kunstleve et al. 2012) until geometry (Ramachandran 97.48% allowed, 2.49% favorable) and statistics (R_work_ = 0.2124; R_free_ = 0.1880) were judged acceptable (Table 1). All other crystallographic utilities were found in the PHENIX package (Adams, Afonine et al. 2010). Images of the crystallographic model were traced with PyMOL (Schrödinger, LLC, New York, USA) version 2.0.6. Relative solvent exposure of cysteine residues was calculated with ASAview (Ahmad, Gromiha et al. 2004) and scaled from 0 (no exposure) to 1 (full exposure). Interface analysis was conducted by PISA (EBI) (Krissinel and Henrick 2007). Crystallographic data are registered at the Protein Data Bank under accession code 7B1W.

**Table 1.** Crystallographic data collection and model refinement statistics.

|  |  |
| --- | --- |
|  | CrRPE1 7b1w |
| Wavelength (Å) | 0.9679 |
| Resolution range (Å) | 49.29 - 1.935 (2.005 - 1.935) |
| Space group | P 1 21 1 |
| Unit cell (Å, °) | 77.293 297.322 77.747<br>90 116.301 90 |
| Total reflections | 1650136 (151270) |
| Unique reflections | 229414 (21621) |
| Multiplicity | 7.2 (7.0) |
| Completeness (%) | 98.65 (93.36) |
| Mean I/sigma(I) | 9.79 (1.40) |
| Wilson B-factor | 30.34 |
| R-merge | 0.1273 (1.049) |
| R-meas | 0.1372 (1.131) |
| R-pim | 0.05063 (0.4178) |
| CC1/2 | 0.997 (0.616) |
| CC* | 0.999 (0.873) |
| Reflections used in refinement | 229338 (21617) |
| Reflections used for R-free | 2409 (227) |
| R-work | 0.1880 (0.2890) |
| R-free | 0.2124 (0.3261) |
| CC(work) | 0.966 (0.744) |
| CC(free) | 0.961 (0.633) |
| Number of non-hydrogen atoms | 22618 |
| macromolecules | 20867 |
| ligands | 12 |
| solvent | 1739 |
| Protein residues | 2772 |
| RMS(bonds) | 0.008 |
| RMS(angles) | 0.90 |
| Ramachandran favored (%) | 97.48 |
| Ramachandran allowed (%) | 2.49 |
| Ramachandran outliers (%) | 0.04 |
| Rotamer outliers (%) | 0.52 |
| Clash score | 6.26 |
| Average B-factor | 34.73 |
| macromolecules | 34.24 |
| ligands | 34.16 |
| solvent | 40.61 |

### Small angle X-rays scattering

50 μL of pure CrRPE1 concentrated at 9.5 mg mL^−1^ were injected on BioSEC-3 300 size-exclusion chromatography column (Agilent Technologies, Santa Clara, USA) equilibrated in buffer 20 mM Tris-HCl (pH 7.9) and 100 mM NaCl, in line with the small-angle X-ray scattering (SAXS) exposure capillary at the synchrotron beamline SWING (SOLEIL, Saint Aubin, France). Collected diffusion images were analyzed on the application Foxtrot 3.3.4 (Xenocs, Sassenage, France) and the ATSAS 2.8.3 suite (Petoukhov, Franke et al. 2012, Franke, Petoukhov et al. 2017). PrimusQT calculated a radius of gyration of 35.49 ± 0.03 Å and estimated SAXS CrRPE1 molecular weight at 173,354 Da (Qp), 169,124 Da (MoW), 158,056 Da (Vc), 165,791 Da (size and shape). Oligomeric state was finally computed as mean SAXS molecular weight of 166,581 Da, divided by monomeric recombinant CrRPE1 sequence molecular weight of 26,773 Da and resulting in an estimated number of 6 subunits per oligomer.

### Algal protein extracts preparation

A single colony of wild-type *Chlamydomonas reinhardtii* strain D66 was grown in 50 mL TAP medium (Gorman and Levine 1965) under continuous 120 rpm agitation and illumination at an intensity of 36 μE s^−1^ m^−2^. At 19.6*10^6^ cells mL^−1^ the culture was harvested by 5 min centrifugation at 3,000 *g* and stored at −20 °C. Cell pellet (600 mg) was thawed and resuspended in 12 mL of 20 mM Tris-HCl supplemented with 100 mM NaCl (pH 7.9), and lysed by passage through CellD (Constant systems, Daventry, UK) at 30 kPSI. Soluble fraction was separated by 10 min centrifugation at 30,000 *g* and filtered through 0.2 μm membrane. The resulting suspension (300 μL) was injected on a Superose 6 Increase 10/300 GL column and eluted isocratically at 0.3 mL min^−1^ in 20 mM Tris-HCl supplemented with 150 mM KCl (pH 7.5). Elution fractions (400 μL) were collected, starting from column dead volume 7.2 mL (fraction 18) and up to column total volume 28.0 mL (fraction 70), snap-frozen in liquid nitrogen, and stored at −20 °C for western blot detection using rabbit polyclonal antibodies raised against recombinant CrRPE1 (Covalab, Bron, France).

### Western blot

Size-exclusion chromatography of Chlamydomonas soluble protein extract fractions 19 to 44 were mixed with reducing Laemmli loading buffer, boiled and resolved by 200 V electrophoresis through 12% SDS-PAGE (Weber and Osborn 1969). Proteins were subsequently transferred to Protran 0.2 μm nitrocellulose membrane (GE Healthcare, Chicago, USA). Efficiency of protein transfer was checked by Ponceau staining. Primary polyclonal anti-CrRPE1 antibody was incubated overnight at a dilution of 1:5,000. Secondary anti-rabbit IgG coupled to peroxidase (Sigma-Aldrich reference A9169, Saint Louis, USA) was incubated 2 h at a dilution of 1:10,000 and revealed by ECL Prime colorimetric assay (GE Healthcare, Chicago, USA).

### Circular dichroism spectroscopy

Samples of CrRPE1 (10.7 μM) were prepared in Tris-HCl buffer (30 mM, pH 7.9) and quantified by spectrophotometric analysis at 280 nm in a 1 cm cell (ε_280_ = 13980 M^−1^ cm^−1^ based on primary structure) (Pace, Vajdos et al. 1995). Far-UV circular dichroism (CD) spectra (250–195 nm) were measured at room temperature on a J-810 spectropolarimeter (Jasco, Japan), using a QS-quartz cell with 0.5 mm optical pathlength (Hellma Analytics, Germany), a 2 nm spectral bandwidth, a 20 nm min^−1^ scanning speed, a 4 sec data integration time, a 0.2 nm data interval and an accumulation cycle of 3 scans per spectrum. The resulting CD spectra were blank-corrected and converted to molar units per residue (Δε_res_, in M^−1^ cm^−1^).

## Results

### Quaternary structure of recombinant and native CrRPE1

Based on previous findings, chloroplast RPE from land plants assembles in a homohexameric structure (Kopp, Kopriva et al. 1999, Kopriva, Koprivova et al. 2000, Jelakovic, Kopriva et al. 2003), albeit an octameric form was initially assigned (Chen, Hartman et al. 1998, Teige, Melzer et al. 1998). The cytosolic counterpart, however, was incontrovertibly found as a dimer (Karmali, Drake et al. 1983, Kopriva, Koprivova et al. 2000). To assess the oligomerization state of CrRPE1, the enzyme was heterologously expressed, purified to homogeneity, and analyzed by size exclusion chromatography (SEC, Figure S3A). We observed a single monodisperse peak with an apparent molecular weight of 134 kDa, which corresponds to roughly five times that of a 27 kDa subunit. The oligomeric nature of the protein along with a precise estimate of the number of subunits was further investigated by Small Angle X-ray Scattering (SAXS) (Figures S3B and S3C). The X-ray diffusion curve was analyzed by ATSAS PRIMUS-QT which derived a radius of gyration of 35.53 Å and a molecular weight of 166,581 Da corresponding to six subunits per particle. Because SAXS is considered a more rigorous method to estimate protein molecular weights with respect to analytical chromatography, we considered that CrRPE1 folds as a homohexamer.

To further inspect the oligomerization state of CrRPE, we analyzed the quaternary structure of the native enzyme extracted from Chlamydomonas cell cultures. A clone of Chlamydomonas D66 strain was grown under constant illumination and after cell lysis, we submitted the soluble fraction to analytical SEC (Figure S3D). Elution fractions were resolved by denaturing electrophoresis (SDS-PAGE), transferred to nitrocellulose, and immunoblotted using primary polyclonal anti-CrRPE1 antibodies. Fractions containing detectable amounts of RPE corresponded to elution volumes comprised between 15.6 mL and 17.2 mL, with a maximum signal in the 16.0-16.4 mL fraction (Figure S3E). This elution volume matches that of the recombinant protein (16.25 mL, Figure S3A) that we observed on the same chromatographic system. We conclude that RPE1 extracted from cultivated algae has the same molecular weight as the pure recombinant protein, and that CrRPE1 assembles as an homohexamer *in vivo*.

### CrRPE1 folds as an α8β8 triose-phosphate isomerase (TIM) barrel

We solved the three-dimensional structure of CrRPE1 by X-ray crystallography at a resolution of 1.9 Å. CrRPE1 crystallized in the monoclinic space group P 1 2_1_ 1 with twelve subunits that assemble in hexameric structures (Figure 1). Crystal packing analysis thus confirmed the homohexameric organization of CrRPE1, in agreement with the quaternary structure assessed by SAXS analysis. All subunits are virtually identical (RMSD = 0.122-0.171 Å) and in the representative chain A, residues Thr31 to Pro260 of mature protein were modelled into continuous electron density. Each CrRPE1 monomer adopts a canonical (β/α)_8_-barrel fold (TIM-barrel, CATH topology 3.20.20, SCOPe family c.1.2.2) with 8 parallel β-strands forming the central cylindrical β-sheet surrounded by 8 α-helices joined by loops (Figure 2). An additional short amino-terminal α-helix caps the basal, amino-exposing side of the β-barrel. The overall content of secondary elements in the crystal structure (helices: 28.1%; strands: 17.4%; turns: 12.2%; other: 42.3%) perfectly reflects the TIM-barrel fold and this was further confirmed by in-solution analysis using CD spectroscopy (Figure S4).

**Figure 1.**
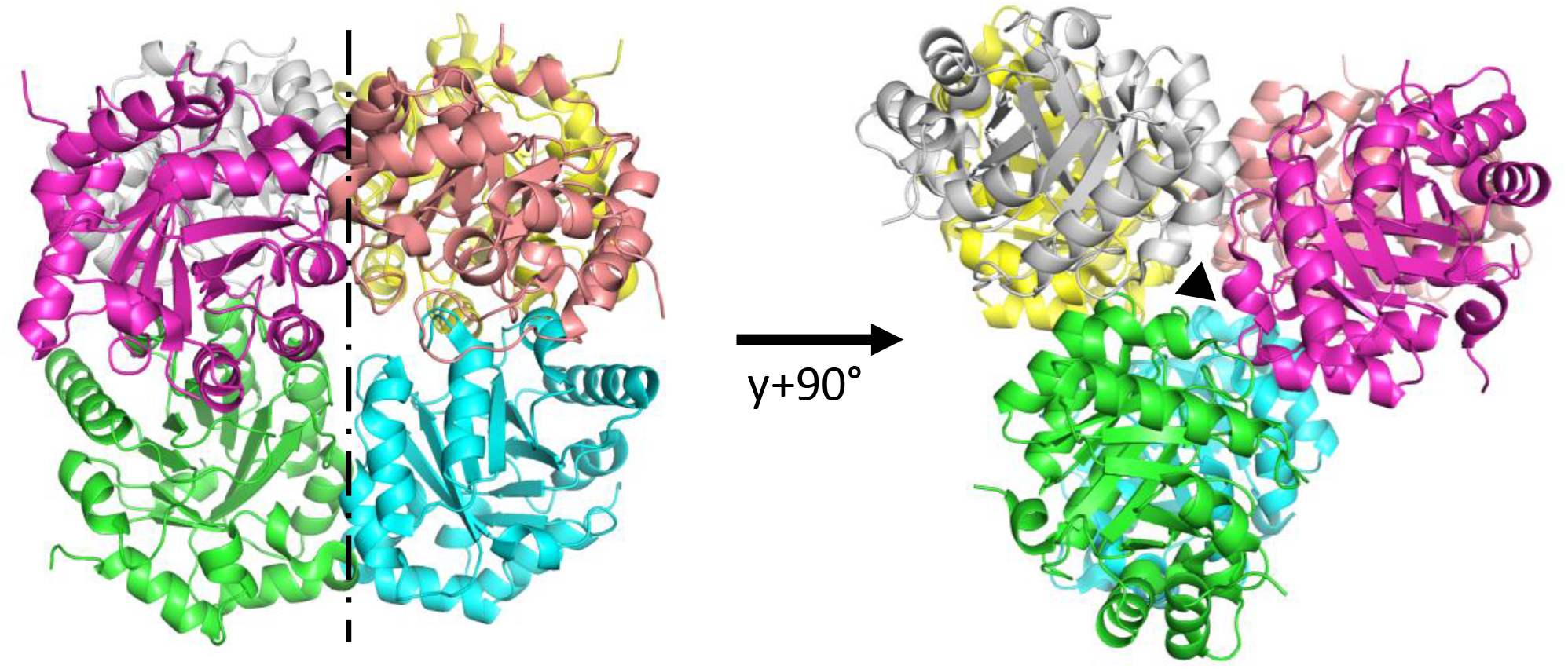
Crystal structure of ribulose-5-phosphate 3-epimerase isoform 1 from *Chlamydomonas reinhardtii* (CrRPE1). Quaternary structure of CrRPE1. Subunits of the homo-hexamer are colored. Projected views highlight the dimerization axis on the left, and the trimerization axis on the right.

**Figure 2.**
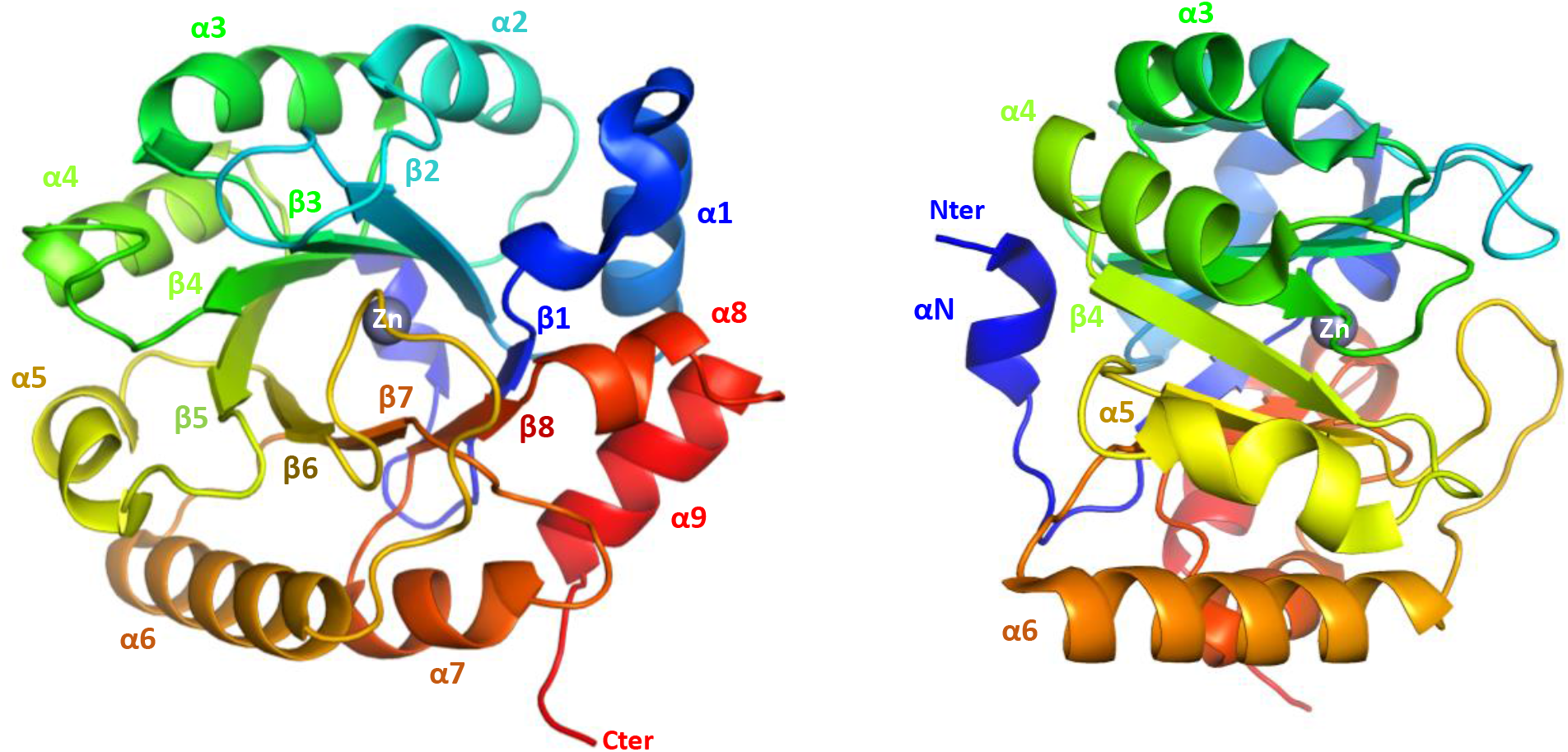
CrRPE1 folding. Main chain is traced in cartoon colored from blue (amino-terminus) to red (carboxy-terminus). Secondary structure elements are annotated. A zinc ion is placed in the center top of the β-barrel.

### CrRPE1 packs as a trimer of dimers

Crystal packing analysis identified nearest neighbors in and around the unit cell. EBI-PISA most probable assembly is composed of the six chains ABDFGK, burying a total surface of contact of 14,140 Å^2^. Computed free energy of dissociation is 20.3 kJ.mol^−1^. Three equivalent interfaces form sub-dimers between chains G-A, K-D, and F-B while six additional interfaces form between B-A, D-B, D-A, K-G, G-F, and K-F to trimerize the dimers into the full hexamer (Figure 1). In the representative G-A dimer, the interface involves ten hydrogen bonds: Leu56_G_-Ser54_A_, Gly57_G_-Ser54_A_, Gly87_G_-Thr85_A_, Arg79_G_-Glu110_A_ (two bonds), Ser54_G_-Leu56_A_, Ser54_G_-Gly57_A_, Thr85_G_-Gly87 _A_, Glu110_G_-Arg79_A_ (two bonds), and five salt bridges: Arg79_G_-GIu110_A_ (three bridges), and the mirror equivalent residues pair Glu110_G_-Arg79_A_ (two bridges). The interface significance score is 1, implying it as essential for complex formation. Trimer interfaces such as that of subunits B-A only form with six hydrogen bonds, in residue pairs Ser133_B_-Gln132_A_ (two bonds), Ser161_B_-Glu166_A_, Phe80_B_-His137_A_, Asp181_B_-Arg140_A_ and the mirror pair Ser163_B_-Ser163_A_. Complex significance score is 0.328, attributing an auxiliary role to these polar interactions in the stabilization of the full complex. The assembly sequence for hexameric RPE probably starts with the association of dimers, before they can contribute to several weak trimerization contacts that eventually gather the full hexamer altogether. This oligomeric organization was previously described for RPE isoforms from *Solanum tuberosum, Synechocystis*, and *Streptococcus pyogenes* (Figure S5) (Kopp, Kopriva et al. 1999, Wise, Akana et al. 2004, Akana, Fedorov et al. 2006).

### CrRPE1 is a Zinc-containing protein with a conserved catalytic pocket

RPEs from both plant and non-plant sources typically contain a metal ion which has a stabilizing effect towards the substrate (Liang, Ouyang et al. 2011). However, some RPE isoforms with available crystallographic structures do not contain any metal ion highlighting the possibility that RPE folding does not require the presence of a specific metal within the active site. What remains to be established is whether enzymes that lack a metal ion are catalytically functional. Crystal structure analysis of CrRPE1 revealed that a spherical electron density (not corresponding to the protein) is located close to the carboxy-exposing side of the barrel. A zinc ion was modelled into the density (Figure 2), as observed for cytosolic RPEs from *Oryza sativa* (PDB entry: 1H1Z) and from *Streptococcus pyogenes* (PDB entry: 2FLI). The canonical tetrad of residues composed by His72, Asp74, His105 and Asp216 (CrRPE1 numbering) coordinate the metal ion. His72 and Asp74 belong to β-strand 2, while His105 and Asp216 belong to β-strand 3 and β-strand 7, respectively. Besides stabilizing the metal ion, these aspartate residues were attributed a crucial role in the catalytic mechanism exchanging protons with the epimerized carbon atom of the substrate (Chen, Larimer et al. 1999).

In CrRPE1, the zinc ion is positioned at the bottom of a deep cleft sided by loop A (Leu49-Phe53), loop B (Met76-Gly87), and loop C (Val180-Lys188) (Figure 3A). Loops A, B and C respectively project from β-strands 1, 2 and 6. Loops A and B contact each other by hydrophobic bonds between side chains of Leu49 and Thr85, and between side chains of Phe53 and Ile86. Loops B and C make multiple hydrophobic contacts involving the following pairs of residues: Val81-Pro182, Val81-Gly183, Pro82-Gly183, and Pro82-Phe184. The constrained positioning of the three loops restricts the path of an incoming substrate (X5P or Ru5P).

**Figure 3.**
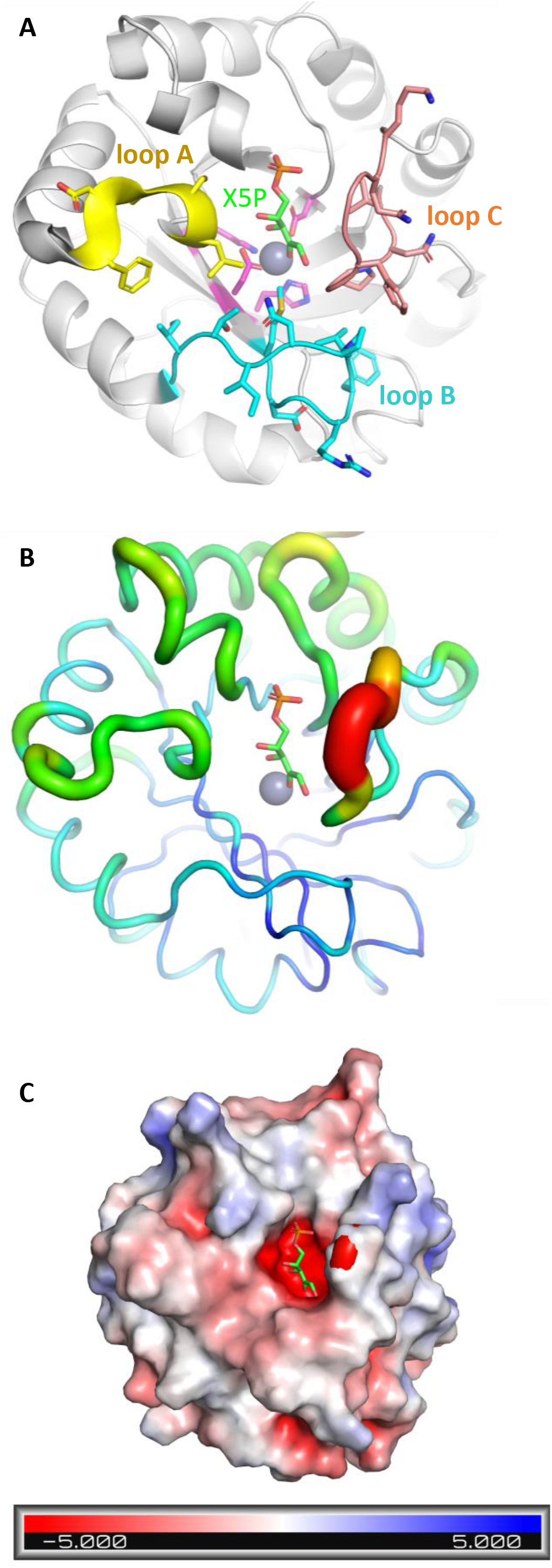
Active site modelization CrRPE1. (A) Cartoon representation of CrRPE1 main chain. Zinc is represented as a gray sphere. Xylulose-5-phosphate (X5P) position was inferred by alignment of CrRPE1 structure with that of *Homo sapiens* RPE co-crystallized with X5P (3OVR (Liang, Ouyang et al. 2011)). Loops A, B, and C surrounding the active site are colored yellow, cyan, and teal with residue side chains represented in sticks. (B) Local mobility of CrRPE1 is represented by crystallographic B-factors: thin, blue ribbon represents low B-factors; large, red ribbon represents high B-factors. (C) CrRPE1 electrostatic surface calculated by PyMOL APBS (Jurrus, Engel et al. 2018) is represented in a gradient from blue (electropositive) to red (electronegative).

In order to predict the substrate binding mode in the active site of CrRPE1, we aligned CrRPE1 crystal structure onto the complex of human RPE (HsRPE) and its substrate X5P (PDB entry: 3OVR) with an RMSD = 0.804 Å. X5P appears positioned in the space defined between the zinc ion and the three aforementioned loops, respecting the lock-and-key complementarity within the active site cleft (Figure 3A). The conformation defined by CrRPE1 specific side chains hence fully accommodates the substrate dimension. The main structural difference between CrRPE1 and HsRPE complexed with X5P is the repositioning of the portion Phe184-Ph189 of the highly conserved loop C closer to the substrate. X5P binding probably induces a fit of the enzyme that closes around its substrate. Accordingly, loop Phe184-Phe189 of our model displays the highest crystallographic B-factors of the whole protein, supporting a functional mobility of these residues upon substrate and product accommodations (Figure 3B). The substrate binding pocket presents an overall electronegative potential (Figure 3C). The 5-phosphatidyl group of the modelled substrate is oriented in proximity to main chain amino groups of Gly217, Gly218, Gly238, and Ser239. Electrostatic interactions between electropositive or hydrogen bond donor amines and negatively charged or hydrogen bond acceptor phosphate favor the alignment of the substrate carbonyl and the hydroxyl group of the C3 substrate at 2.8 Å and 3.4 Å from the zinc ion, respectively. This conformation of the active site appears optimal to ensure efficient X5P to Ru5P interconversions.

### Enzyme kinetic properties of CrRPE

Because large discrepancies appear from kinetic comparison of plant RPE orthologs, we sought to determine the kinetic features of recombinant CrRPE1. To this end, we replicated and optimized protocols from previous studies (Davis, Lee et al. 1972, Kiely, Stuart et al. 1973, Chen, Larimer et al. 1999, Akana, Fedorov et al. 2006). The initial velocity (*v*_i_) of CrRPE1 ascribed to the PPP-related activity (*i.e*., Ru5P epimerization to X5P) was measured by coupling Ru5P epimerization to NADH oxidation via three reporter enzymes (CrTK, CrTPI, α-GDH) (Figure S2). When CrRPE1 was omitted in the assay cuvette, we detected a slow and time-limited NADH consumption likely attributed to X5P contamination of Ru5P powder. Consequently, the further oxidation of NADH observed after adding CrRPE1 in the assay mixture indicates that the formation of X5P by CrRPE1 is strictly required to allow the continuous functioning of coupled enzymes (*i.e*., the CrTK-dependent formation of glyceraldehyde-3-phosphate and subsequent catalysis by CrTPI and α-GDH, Figure S2). After establishing the enzymatic assay, we analyzed the dependency of CrRPE1 activity on protein concentration. As shown in Figure 4A, we found that protein activity (ΔAbs_340_/min) displayed a linear relationship with increasing protein concentration in the 2.5-10 nM range, corresponding to a specific activity of 387.9 ± 30.4 μmol min^−1^ mg^−1^ (Figure S6A). The kinetic parameters were then determined using Ru5P as variable substrate concentration and activity data were analyzed by non-linear regression analysis using the Michaelis-Menten equation (Figure 4B). Enzyme activities plotted versus Ru5P concentration displayed a typical hyperbolic response and CrRPE1 catalyzed the epimerization of Ru5P to X5P with a *K*_M_ value of 1.52 ± 0.19 mM and a turnover number (*k*_cat_) of 272.6 ± 17.4 sec^−1^. By comparing the kinetic properties of CrRPE with recombinant and native RPE from spinach (SoRPE), which is the only plastidial isoform to have been kinetically characterized to date, we noted that the affinity for Ru5P is markedly different as native and recombinant SoRPE showed a ~6-fold lower Michaelis-Menten constant (0.22-0.25 mM) (Table 2). A striking diversity was also observed when comparing turnover numbers since CrRPE catalyzes the reaction with a value about 25-fold lower and 2000-fold higher than those previously reported for the recombinant and native form of spinach RPE, respectively (Table 2) (Chen, Hartman et al. 1998, Teige, Melzer et al. 1998, Chen, Larimer et al. 1999). In view of the values reported here and in previous studies, any comparative conclusion on the catalytic features is hazardous since the catalytic constants for spinach native and recombinant RPE isoforms are extremely discordant. One possible explanation was provided by Chen and coauthors who considered the presence of the reducing agent dithiothreitol (DTT), which was used during the purification procedure of the native form, as a destabilizing agent that alters the functionality of the protein (Chen, Hartman et al. 1998). To further inspect the kinetic diversity with SoRPE, the spinach enzyme was recombinantly expressed, purified, and biochemically characterized (Figure 4A). Surprisingly, the recombinant SoRPE catalyzed the epimerization of Ru5P to X5P with a specific activity ~2-fold lower compared to CrRPE (Figure S6A) and exhibited a similar *K*_M_ with respect to CrRPE1 (1.56 ± 0.17 mM) and 2.5-fold lower *k*_cat_ value (105.4 ± 13.5 sec^−1^) (Figure S6B). Taken together, our results on spinach enzyme are in sharp contrast to previous studies differing for both substrate affinity and maximal activities (*i.e*., specific activity or turnover numbers) (Table 2). In this regard, several reasons can be proposed that may depend on multiple factors: (i) the functional and redox state of the RPE enzyme, (ii) the source and specificity of the coupled enzymes, and (iii) the purity and integrity of the chemicals used. However, other possible explanations could justify these differences, and for this reason we do not intend to question further this striking discrepancy.

**Figure 4.**
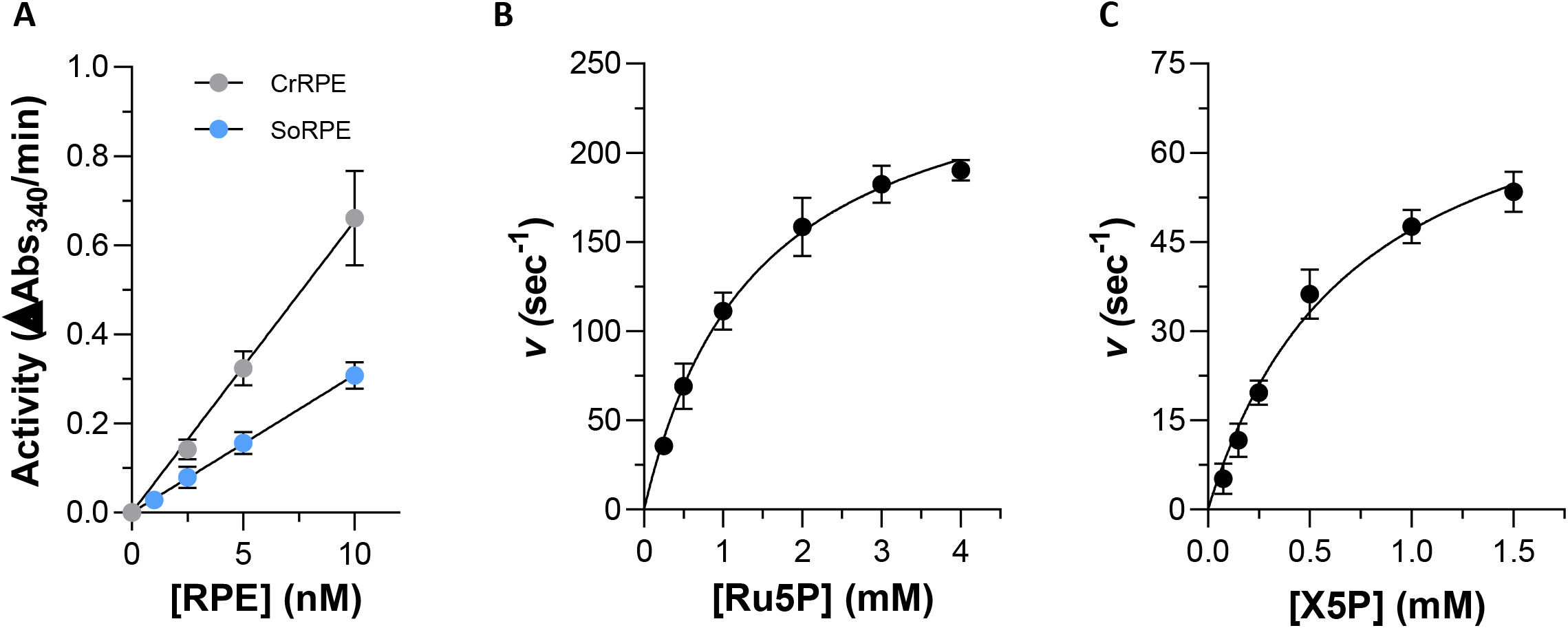
*In vitro* enzymatic activity of *Chlamydomonas reinhardtii* (Cr) RPE and of *Spinacia oleracea* (So) RPE. (A) Assay linearity in a 0-10 nM range of enzyme concentration. (B) Ribulose-5-phosphate (Ru5P) saturation function in a 0-4 mM range of substrate concentration. (C) Xylulose-5-phosphate (X5P) saturation function in a 0-1.5 mM range of substrate concentration.

**Table 2.**
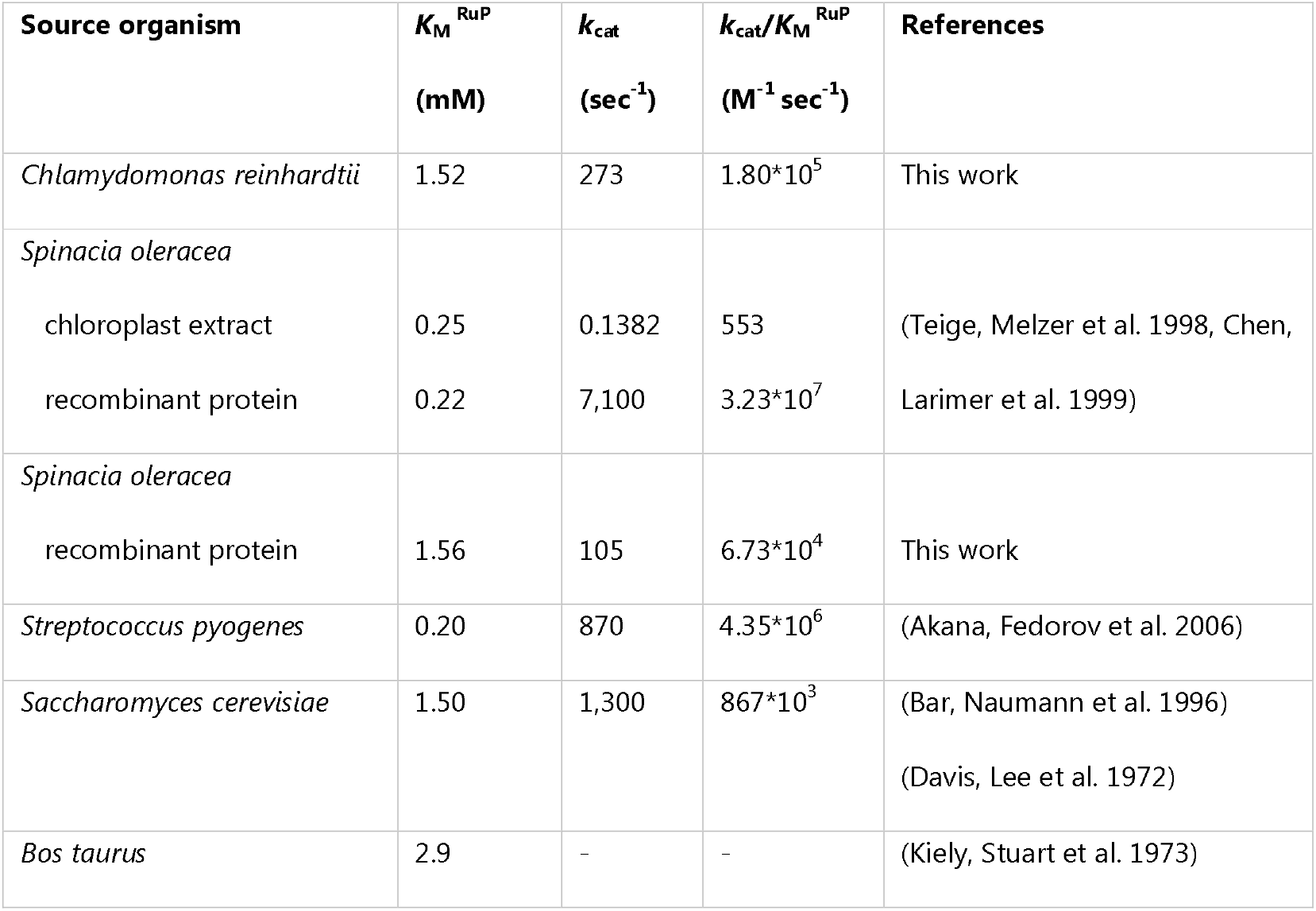
Kinetic parameters of RPE. Catalytic parameters of RPE from *Chlamydomonas reinhardtii* and *Spinacia oleracea* (this study), and (Teige, Melzer et al. 1998, Chen, Larimer et al. 1999), *Streptococcus pyogenes* (Akana, Fedorov et al. 2006), *Saccharomyces cerevisiae* (Davis, Lee et al. 1972, Bar, Naumann et al. 1996), and *Bos taurus* (*Kiely, Stuart et al. 1973*).

| Source organism | $K_M^{\text{RuP}}$<br>(mM) | $k_{\text{cat}}$<br>($\text{sec}^{-1}$) | $k_{\text{cat}}/K_M^{\text{RuP}}$<br>($\text{M}^{-1} \text{sec}^{-1}$) | References |
| --- | --- | --- | --- | --- |
| *Chlamydomonas reinhardtii* | 1.52 | 273 | $1.80 \times 10^5$ | This work |
| *Spinacia oleracea* |  |  |  |  |
| chloroplast extract | 0.25 | 0.1382 | 553 | (Teige, Melzer et al. 1998, Chen, |
| recombinant protein | 0.22 | 7,100 | $3.23 \times 10^7$ | Larimer et al. 1999) |
| *Spinacia oleracea* |  |  |  |  |
| recombinant protein | 1.56 | 105 | $6.73 \times 10^4$ | This work |
| *Streptococcus pyogenes* | 0.20 | 870 | $4.35 \times 10^6$ | (Akana, Fedorov et al. 2006) |
| *Saccharomyces cerevisiae* | 1.50 | 1,300 | $867 \times 10^3$ | (Bar, Naumann et al. 1996) |
|  |  |  |  | (Davis, Lee et al. 1972) |
| *Bos taurus* | 2.9 | - | - | (Kiely, Stuart et al. 1973) |

Whereas PPP-related RPE activity has been widely employed to dissect the catalytic properties of RPE enzymes, there is a lack of knowledge about the biochemical features of RPE related to the conversion of X5P into Ru5P (i.e., CBBC-related activity). To assess the catalytic properties related to the catalytic capacity to use X5P as substrate, we employed the enzymatic assay typically used to monitor the activity of phosphoribulokinase (PRK) (Gurrieri, Del Giudice et al. 2019). The enzyme catalyzes the ATP-dependent phosphorylation of Ru5P, and in the assay we developed, the substrate of PRK is provided by CrRPE1 through epimerization of X5P. To this respect, we observed that the CrRPE1 amount in the assay cuvette was the only limiting factor of the assay. Consistently, we found that protein activity (ΔAbs_340_/min) displayed a linear relationship with increasing protein concentration in the 2.5-20 nM range (data not shown). The derived specific activity was 115.7 ± 6.7 μmol min^−1^ mg^−1^, which is ~3.3-fold lower compared to the PPP-related activity (Figure S6A). Kinetic constants were then determined by using variable X5P concentrations (0-1.5 mM) and activity data were analyzed by non-linear regression analysis using the Michaelis-Menten equation (Figure 4C). Enzyme activities plotted versus X5P concentration displayed a typical hyperbolic response and CrRPE1 catalyzed the epimerization of X5P to Ru5P with a *K*_M_ value of 0.716 ± 0.09 mM and a *k*_cat_ of 80.7 ± 7.9 sec^□1^. Overall, these results indicate that CrRPE1 has a higher affinity for X5P compared to Ru5P but displays a lower capacity to convert X5P into Ru5P with respect to the reverse reaction. Consequently, derived catalytic efficiencies (*k_cat_/K*_M_) slightly differ from 1.13 × 10^5^ M^−1^ sec^−1^ for the enzyme activity employing X5P as a substrate to 1.80 × 10^5^ M^−1^ sec^−1^ for the epimerization of Ru5P to X5P.

### CrRPE1 displays a limited intrinsic instability and redox sensitivity to reducing agents

The stability of a given protein under different cellular conditions (*e.g*., low temperature or reducing conditions) is a crucial parameter because of the direct connection between structural integrity and functionality. In a previous study, Chen and colleagues showed that the chloroplast SoRPE (recombinant form) appeared to be highly unstable even under control conditions at 4 °C (Chen, Hartman et al. 1998). Moreover, the activity of the recombinant enzyme was strongly affected after exposure to reducing agents such as 2-mercaptoethanol. In relation to this evidence, we investigated the response of CrRPE1 activity to incubation at low temperature (4 °C) and treatments with two chemical reducing agents, namely 2-mercaptoethanol and dithiothreitol (DTT). CrRPE1 activity was totally unresponsive to temperature (~99% activity compared to control) whereas a mild reduction in activity was observed following exposure with 2-mercaptoethanol (~40% inhibition after 2 h incubation). Intriguingly, no significant alteration of the enzymatic activity was detected upon treatment with DTT. Although DTT and 2-mercaptoethanol share similar reactivity toward protein cysteines, the redox mechanism underlying the inhibitory effect of the latter is unclear. In this regard it is worth noting that a reductive-based redox mechanism would imply a pre-existing oxidized state of the protein, which would be subsequently modified by reducing treatments. However, this is in contrast with the observation that in the crystal structure all cysteine thiols are found in a reduced state (Figure 5A). Therefore, we can assume that the observed inactivation mediated by 2-mercaptoethanol eludes a cysteine-based redox-type mechanism but rather derives from other properties of the molecule likely affecting protein folding and/or metal coordination.

**Figure 5.**
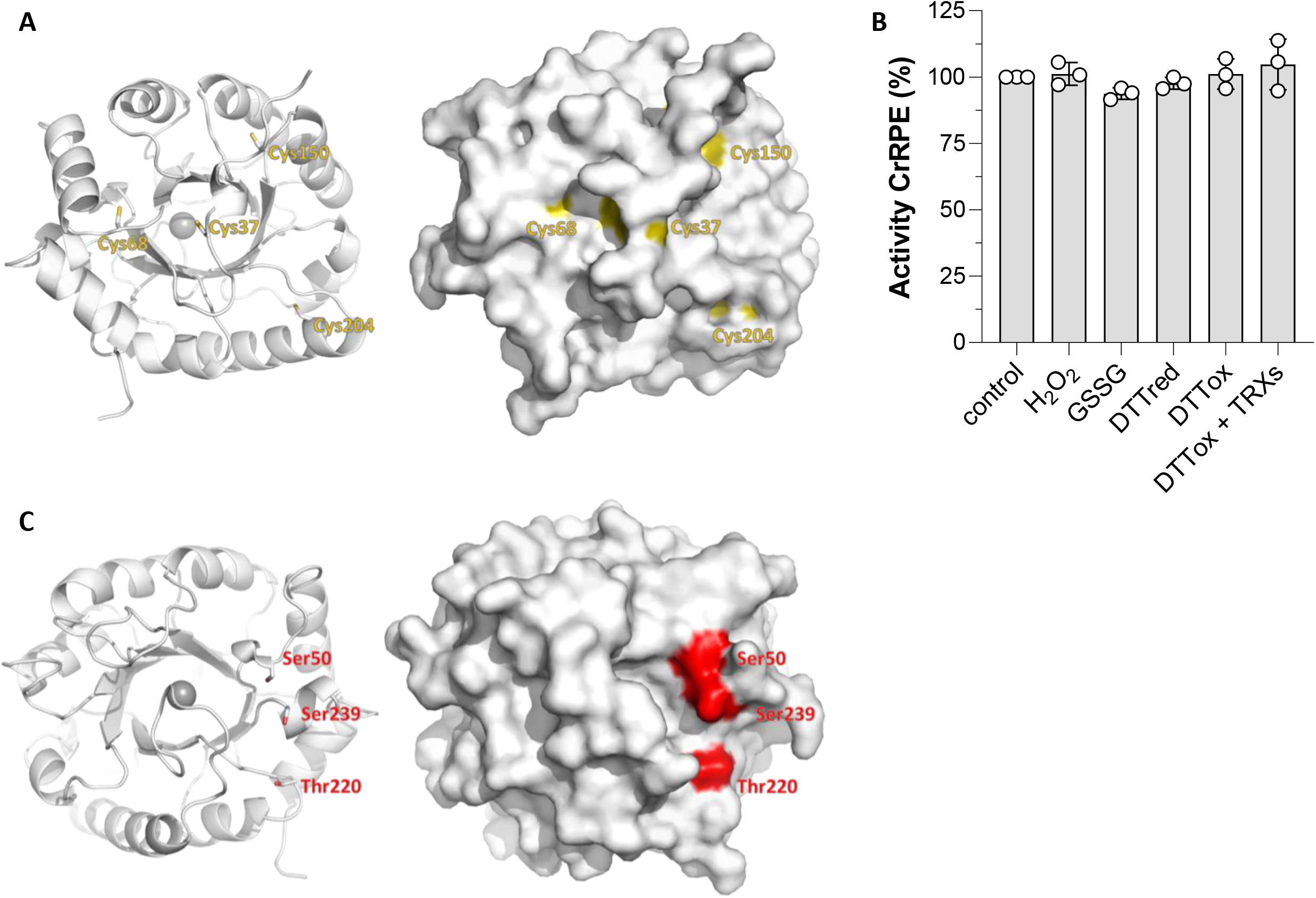
Post-translational modification sites of CrRPE1. (A) Left, cartoon representation of the protein main chain. Potential redox sites identified in Chlamydomonas protein extracts. Right, surface representation with cysteines side chains shown in sticks. Potential redox sites are painted in gold. (B) In vitro enzymatic activity of recombinant RPE submitted to redox treatments. (C) Phosphorylation sites identified in Chlamydomonas protein extracts. Left, cartoon representation of the protein main chain with target amino acids side chains represented in sticks. Right, surface representation. Phosphorylation sites are painted in red.

### Redox response of CrRPE1 to oxidative treatments

CrRPE1 was identified as putative target of TRX implying that it might contain cysteine residues involved in the formation of one or more disulfide bonds (Pérez-Pérez, Mauriès et al. 2017). Besides, CrRPE1 was also found to undergo S-nitrosylation (Morisse, Zaffagnini et al. 2014) and S-glutathionylation (Zaffagnini, Bedhomme et al. 2012). Our crystal structure of CrRPE1 positions the four cysteines with relative surface accessibility of 0.180 for Cys37, 0.076 for Cys150, and 0.000 for both Cys68 and Cys204 (Figure 5A). The thiol of Cys37 is the most accessible to the solvent and therefore it is likely able to react with oxidative molecules, even though minor conformational changes may allow the other cysteines to partially or fully expose their side chains to the solvent. To determine the number of accessible cysteine thiols, we employed the Ellman’s reagent (5,5-dithio-bis-(2-nitrobenzoic acid), DTNB) and measured one accessible thiol group *in vitro* (1.0 ± 0.2), suggesting that only one cysteine is likely competent for redox exchange.

We then examined the redox sensitivity of CrRPE1 after exposure with oxidized DTT alone or in combination with pure recombinant chloroplast thioredoxins from Chlamydomonas. As shown in Figure 5B, no significant alteration of protein activity was observed indicating that dithiol/disulfide interchanges do not constitute a regulatory mechanism of CrRPE1 activity. The catalytic response of CrRPE1 to oxidative modifications was also assessed in the presence of nitrosoglutathione (GSNO), oxidized glutathione (GSSG), and hydrogen peroxide (H_2_O_2_) alone or in combination with reduced glutathione (GSH). Again, we did not observe any variation of CrRPE1 activity with respect to control conditions (Figure 5B). Taken together, these results led us to the conclusion that CrRPE1 catalysis is insensitive to possible variations in the redox state of the cysteine residues, and this correlates with the distal positions of the cysteines from the active site (Figure 5A). Alternatively, we can hypothesize that redox post-translational modifications may have non-catalytic functions, possibly causing local conformational changes that influence interactions with possible partner proteins, or alternatively inducing moonlighting functions whose role is as yet unknown.

### Post-translational phosphorylation sites

Phosphorylation sites Ser50, Thr220 and Ser239 revealed by proteomics (Wang, Gau et al. 2014) map at the vicinity of the active site, at respectively 11 Å, 17 Å and 13 Å of the zinc ion. In our crystal structure, the three residues expose to solvent their unmodified hydroxyl side chains in a 14 Å arch that aligns at the entrance of the active site (Figure 5C). Possible recognition and modification by a kinase are allowed by this solvent exposure. RPE substrates (*i.e*., X5P and Ru5P) are expected to be exchanged with solvent following a path gated by Ser50, Thr220 and Ser239. Phosphorylation of one or more of these sites may hence hinder the formation of the Michaelis-Menten complex, reducing the activity of the enzyme. To assess this hypothesis, we designed and prepared three mutated proteins in which serine or threonine were substituted by aspartates to tentatively mimic with β-carboxylates the introduction of phosphate charges. However, the activity of phospho-mimicking single mutants was not significantly reduced as compared to wild-type (data not shown) either because phosphorylation introduces more charges and larger steric hindrance than the carboxylate mimics, or because more than one site needs to be modified to affect the enzymatic activity.

## Discussion

In this study we describe the novel structure of RPE1 from the model microalga *Chlamydomonas reinhardtii* determined by X-ray crystallography at a resolution of 1.9 Å. Recombinant CrRPE1 purifies as a homo-hexamer built from the trimerization of dimers. We confirmed that the homo-hexamer is the main population in solution and that native RPE extracted from algae cultures also elutes at the same apparent molecular mass as the purified recombinant protein. Hexamerization is not observed in the crystal structure of non-plant RPE (Liang, Ouyang et al. 2011) but is reported for another photosynthetic RPE (Kopp, Kopriva et al. 1999). We observe in our model that the catalytic pocket lies close to both the dimerization and the trimerization interfaces. We propose that the formation of homo-oligomers of RPE supports allosteric cross-talk between subunit active sites, possibly by restricting the mobility of loops around the catalytic pocket. Interestingly, this could contribute to the variation of the enzymatic parameters reported for photosynthetic and non-photosynthetic RPE. The 3D-structure computes B-factors that quantify the local mobility of the protein in the crystal. We observe four elements of high mobility surrounding the active site (Figure 3B), that are good candidates for such an allosteric regulation. The determination of RPE high-resolution structures in less constrained environments, *e.g*., by nuclear magnetic resonance (NMR) or single-particle cryogenic electron microscopy (cryo-EM), would provide more information on the dynamic around the active site.

RPE is a ubiquitous enzyme with conserved active site elements and participates to either PPP or CBBC catalyzing the configuration exchange between Ru5P and X5P. CrRPE1 is a zinc-containing enzyme localized in the chloroplast stroma and it likely performs both epimerization reactions given that both pathways co-exist in the same cell compartment. Catalytic interconversion between X5P and Ru5P consists in an acid-base catalysis involving proton abstraction and donation from and to the C3 atom of the substrate and is proposed to occur through the formation of a 2-3-ene-diolate intermediate (Chen, Larimer et al. 1999, Akana, Fedorov et al. 2006). Active site zinc ion along with conserved methionine residues (Met76, Met107, and Met178) participate in the stabilization of this high energy intermediate interacting with carbon 2 and carbon 3 hydroxyl oxygens and carbon 2 oxyanion. Asp74 and Asp216 successively complete the metal coordination sphere and can exchange protons with the reaction intermediates. Depending on the bound substrate, the reaction proceeds with its deprotonation being carried out by one of the two catalytic Asp (i.e., Ru5P to X5P conversion) or with the taking of a proton from the second catalytic Asp. Based on the kinetic analysis of CrRPE1 here reported, the algal enzyme has similar affinities for X5P and Ru5P with a slightly higher preference for the former. Consequently, the kinetic direction of the reaction could solely depend on the intracellular concentration of X5P/Ru5P owing to the similar tendency of the two different substrates to bind.

While the enzymatic mechanism is clearly established for several RPE orthologs, it is not yet known whether the enzyme undergoes any regulatory mechanisms. In the present study, we mapped for the first time the sites of post-translational modifications that may contribute to the modulation of RPE activity according to the environmental conditions or the energetic demand of the cell. Recent proteomic studies on Chlamydomonas cell extracts revealed that RPE has three sites of phosphorylation (Ser50, Thr220, and Ser239) (Wang, Gau et al. 2014, McConnell, Werth et al. 2018). Notwithstanding their distant position in the sequence, all three residues are located in a small area in close proximity to the catalytic pocket. We propose that insertion of one, two or three bulky charged phosphates might interfere with the binding of the X5P/Ru5P substrate and Ru5P/X5P product release, thus slowing down RPE catalysis. Phosphorylation sites (Ser50, Thr182, and Ser239) are variably conserved in other RPE isoforms from photosynthetic organisms and also in the human enzyme, suggesting that phosphorylation might constitute a conserved regulation mode of RPE activity. Future studies are required to shed light on the possible modulation of photosynthetic-related metabolism as other CBBC enzymes were identified as putative targets of phosphorylation (Wang, Gau et al. 2014, McConnell, Werth et al. 2018).

Cysteine redox status is correlated with photosystems illumination because of the ferredoxin-thioredoxin reduction pathway that was demonstrated to activate several CBBC enzymes by reducing regulatory disulfide bonds (for review see (Michelet, Zaffagnini et al. 2013)). In addition, accessible and reactive protein thiols could act as redox sensors of oxidative stress conditions by reacting with reactive oxygen/nitrogen species resulting primarily in S-nitrosylation or S-glutathionylation of proteins (Zaffagnini, Fermani et al. 2019). CrRPE1 was identified as a putative target of redox modifications, including thioredoxin-mediated dithiol/disulfide exchange (Pérez-Pérez, Mauriès et al. 2017), but also S-nitrosylation and S-glutathionylation (Zaffagnini, Bedhomme et al. 2012, Morisse, Zaffagnini et al. 2014). Based on these premises, we suggest that CrRPE1 enters these redox switches thanks to one or several of its cysteines. Four cysteines are present in the primary sequence of CrRPE1 and thiol titration revealed the presence of a single Cys residue accessible to the solvent. Structural data identified Cys37 as the most accessible to the solvent. However, our *in vitro* attempts revealed that CrRPE1 activity is not modulated by cysteine-based redox modifications, either because the conditions were not stringent enough to induce cysteine oxidation or because the inactive state is unstable. Alternatively, we can consider that cysteine oxidation alone is insufficient to affect protein catalysis and this correlates with the positioning of CrRPE cysteines, which are all located far from the catalytic pocket. Overall, this result is reminiscent of the redox sensitivity of triose-phosphate isomerase (TPI), another CBBC enzyme. This enzyme was identified as a putative redox target, but no significant alteration in protein activity was detected upon exposure to various oxidizing treatments (Zaffagnini, Michelet et al. 2014). Despite being functionally resistant to oxidative modifications, we cannot exclude that alteration of the redox state of CrRPE1 cysteine thiols might induce local conformational changes allowing the enzyme to interact with putative partner proteins. Such interactions would act as a regulatory mechanism as observed for PRK and GAPDH forming an inactive supercomplex bridged by CP12 (Gurrieri, Fermani et al. 2021), or alternatively, they might be functional in the formation of larger enzymatic assemblies to channel CBBC metabolites and increase metabolic efficiency. In this regard, plant RPE was recently proposed to engage in such an activating complex with ribose-phosphate isomerase (RPI) (Kuken, Sommer et al. 2018) and to colocalize with CBBC enzymes in a liquid partition of the stroma (Wang, Patena et al. 2022). Further research on native RPE isolated from algae extracts is needed to evaluate the complexity and dynamics of the RPE proteome and redox-dependent protein-protein interactions.

The current knowledge on kinetic features of plant RPE makes it rather complicated to accurately estimate how the enzyme truly functions in the physiological context of the stroma, which could in turn result into important physiological implications likely to be linked to the substrate to product conversion in the Michaelis-Menten complex. Nonetheless, if we consider the *in vitro* catalytic properties of CrRPE1 related to photosynthetic activity (i.e., CBBC-related activity), we can propose a kinetic modeling in the physiological context of a photosynthetic cell under light conditions. In Chlamydomonas, RPE1 concentration was quantified at 3.3 μM (Hammel, Sommer et al. 2020). Considering that (i) RPE1 is the sole isoform present in the chloroplast stroma with a concentration of 3.3 μM, (ii) the substrate X5P is available at 10-50 μM (Mettler, Muhlhaus et al. 2014), and that (iii) RPE functions with a K_M_ of 0.716 mM and a *k*_cat_ of 81 sec^−1^, we expect an *in vivo* X5P to Ru5P flow ranging from 3.7 to 17 μM sec^−1^. These values are comparable to or lower than those calculated for Rubisco of 18.9 μM sec^−1^ (*k*_cat_ = 5.8 sec^−1^, K_M_= 0.029 mM (Tcherkez, Farquhar et al. 2006), [RuBP] = 200 μM (Mettler, Muhlhaus et al. 2014), RbcS stroma concentration of 183 μM (Hammel, Sommer et al. 2020)). Relying on *in vitro* kinetic constant measurements and metabolome and proteome values of substrates and enzymes concentrations, it is possible to speculate that any increase in RPE protein abundance and optimization of its function may result in improved regeneration efficiency of RuBP and thus increased Rubisco activity as suggested by recent models (Raines 2022). Synthetic biology engineering toolkits available in Chlamydomonas and other microalgae (Li, Zhang et al. 2016, Shin, Lim et al. 2016, Crozet, Navarro et al. 2018) now offer the opportunity to modify the RPE-dependent reaction by either changing protein quantities or altering kinetic features (*i.e*., substrate affinity and turnover number) to improve the catalytic efficiency. Tinkering a synthetic CBBC from engineering principles will both comfort the comprehension of its physico-chemical properties and open important opportunities to improve photosynthetic efficiency and likely crops yield.

## Acknowledgements

This work was funded by CNRS, Sorbonne Université, and Agence Nationale de la Recherche grants LABEX DYNAMO (11-LABX-0011), CALVINDESIGN (ANR-17-CE05-001) and CALVINTERACT (ANR-19-CE11-0009). We acknowledge Marion Hamon and Dr. Christophe Marchand at the mass spectrometry platform of the Institut de Biologie Physico-Chimique (FRC 550 CNRS) for protein identification by MALDI-TOF by peptide mass fingerprinting. The Institut de Biologie Physico-Chimique provided access to crystallization platform. We acknowledge the European Synchrotron Radiation Facility (Grenoble, France) for provision of synchrotron radiation facilities at beamline ID30A-3 MASSIF-3 and SOLEIL (Gif-sur-Yvette, France) for provision of synchrotron radiation facilities on beamlines SWING, Proxima-1 and Proxima-2a.

## Supplementary figures

**Supplementary figure 1. Multiple
 sequence alignment of RPE from photosynthetic eukaryotes.**

Gi|159465721 *Chlamydomonas reinhardtii*, gi|333691285 *Dunaliella salina*, gi|1527553216 *Coffea arabica*, gi|1228842444 *Helianthus annuus*, gi|1226781342 *Spinacia oleracea*,gi|15240250 *Arabidopsis thaliana*, gi|923607646 *Brassica napus*, gi|356511994 *Glycine max*.Secondary structures of CrRPE1 determined from crystallographic model are depicted above alignment. Conserved residues are written in white on a red background. Image was generated with ESPript (Gouet, Courcelle et al. 1999).

**Supplementary figure 2. Schematic view of the enzymatic reporter assays used in this study.**

Calvin-Benson-Bassham cycle epimerization of X5P into Ru5P is reported by PRK, PK, and LDH. Pentose-phosphate pathway related epimerization of Ru5P into X5P is reported by TK, TPI, and α-GDH.

**Supplementary figure 3. *In vitro* quaternary structure analysis of CrRPE1.**

(A) Size-exclusion chromatography profile. Globular standards elution volumes are indicated with molecular weight in kDa. (B) SAXS diffusion curve log(I)=f(s). (C) Distance distribution function calculated from SAXS diffusion curve. (D). Size-exclusion chromatography profile of soluble proteins extracted from Chlamydomonas. (E) Anti-CrRPE1 western blot of chromatography fractions. Elution volumes of the collected fractions (400 μL each) are indicated above the membrane.

**Supplementary figure 4. Recombinant CrRPE1 far-UV circular dichroism spectrum.**

**Supplementary figure 5. Comparison of RPE quaternary structure.**

Homo-hexameric RPE from Chlamydomonas (7b1w, this study), *Synechocystis* (1tqj (Wise, Akana et al. 2004), and *Streptococcus pyogenes* (2fli (Akana, Fedorov et al. 2006)). Two rotated projections are represented with main chains colored differently for each subunit.

**Supplementary figure 6. *In vitro* specific activity of plant RPE.**

Purified recombinant *Chlamydomonas reinhardtii* (Cr) RPE and *Spinacia oleracea* (So) RPE were assayed in the PPP-type reporter setup. (A) Specific activities of CrRPE and SoRPE. (B) Ribulose-5-phosphate (Ru5P) saturation function of SoRPE in a 0-4 mM range of substrate concentration.

