## Supplementary figures and images for "Structural and functional characterization of chloroplast ribulose-5-phosphate-3-epimerase from the model green microalga *Chlamydomonas reinhardtii*"

### Supplementary figures

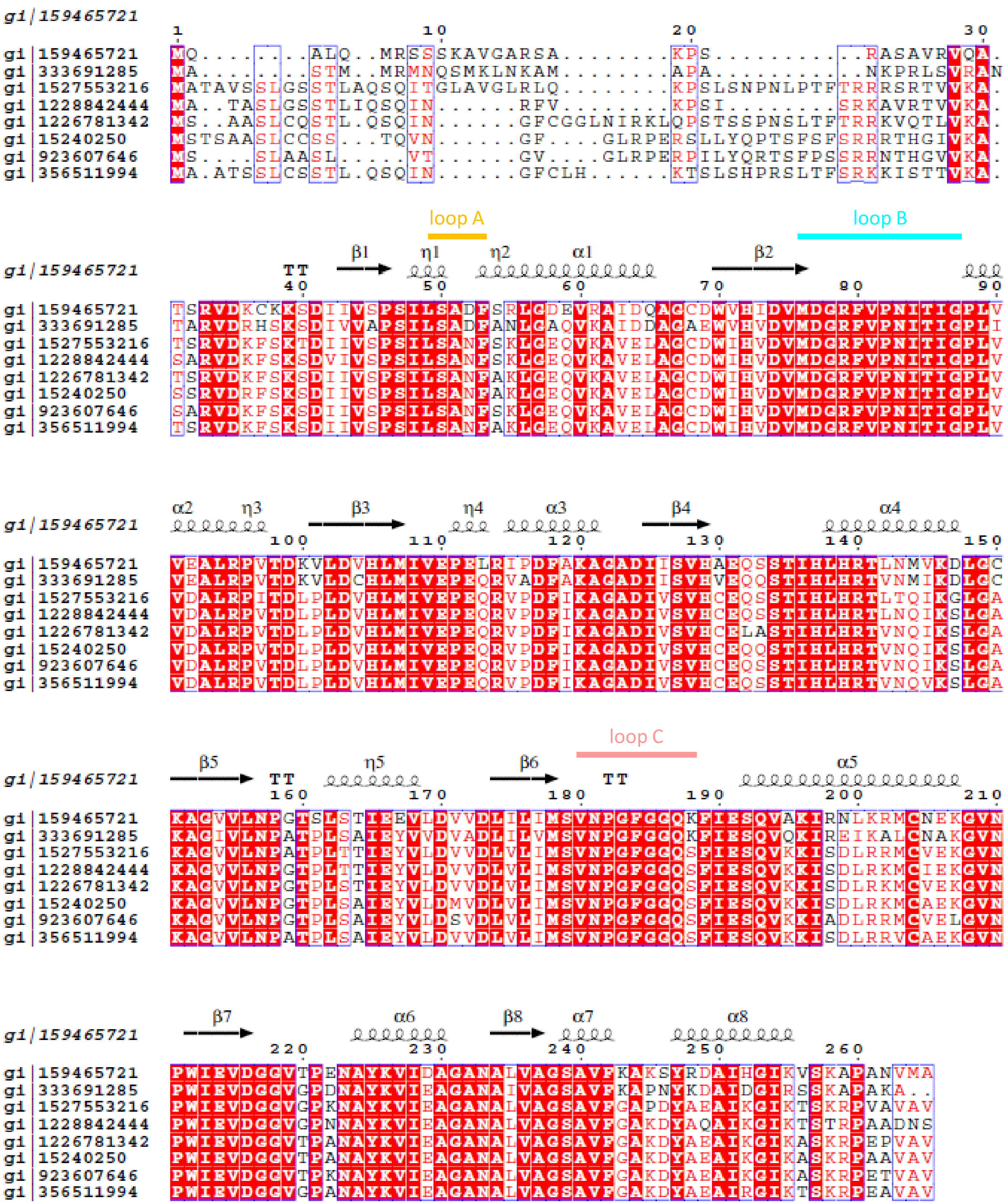

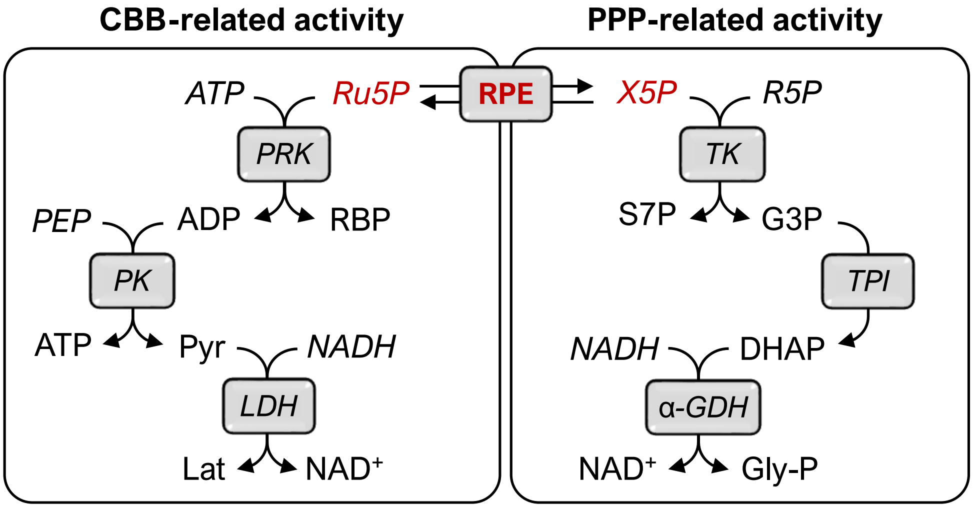

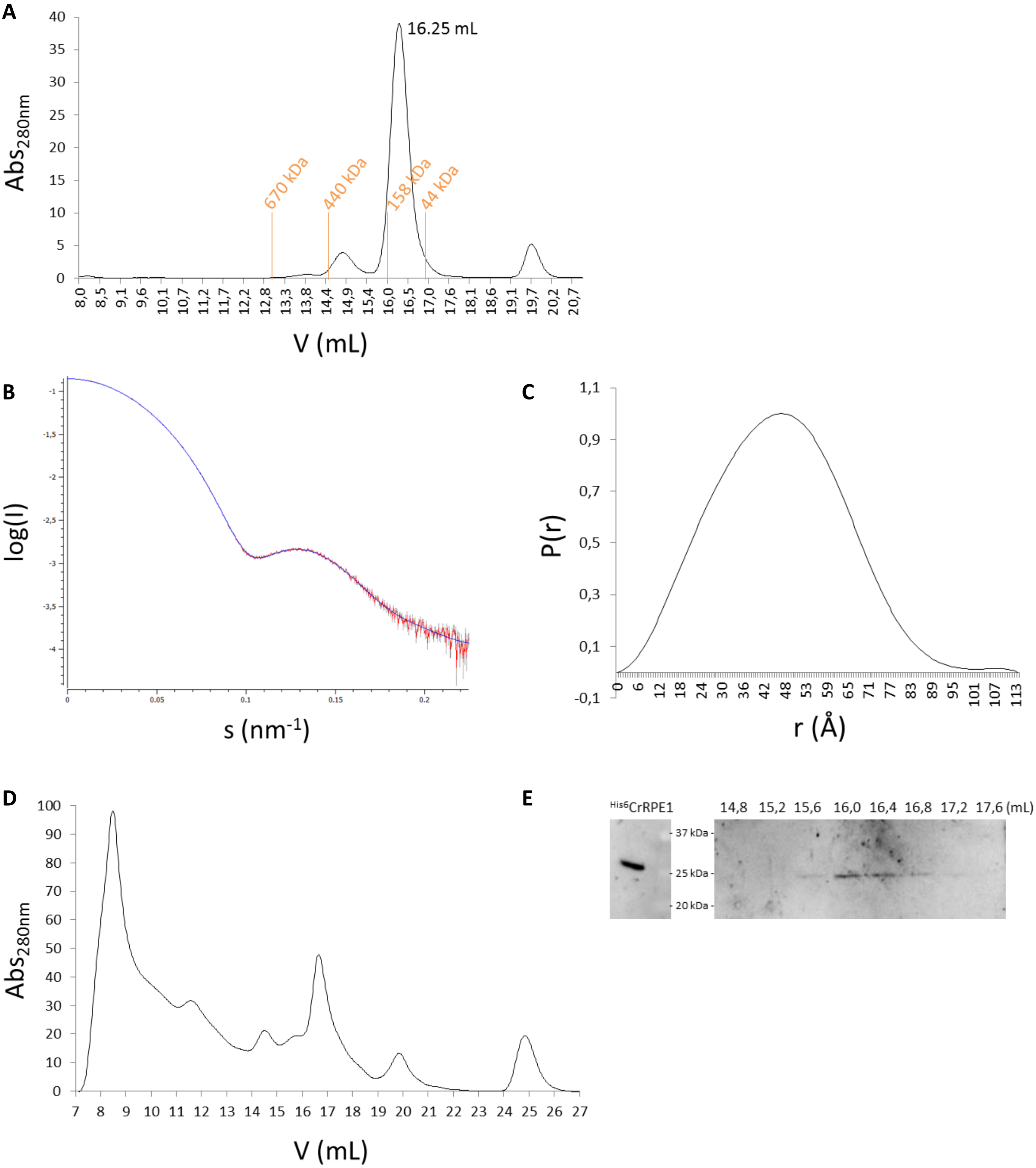

Figure S4

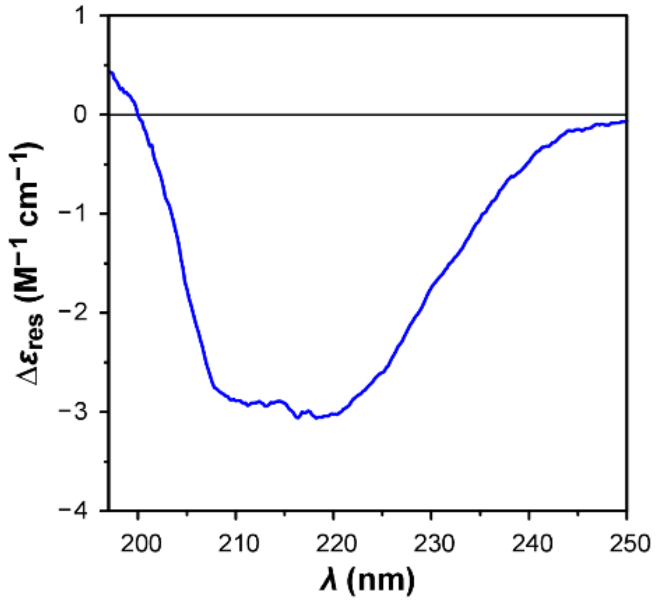

Figure S5

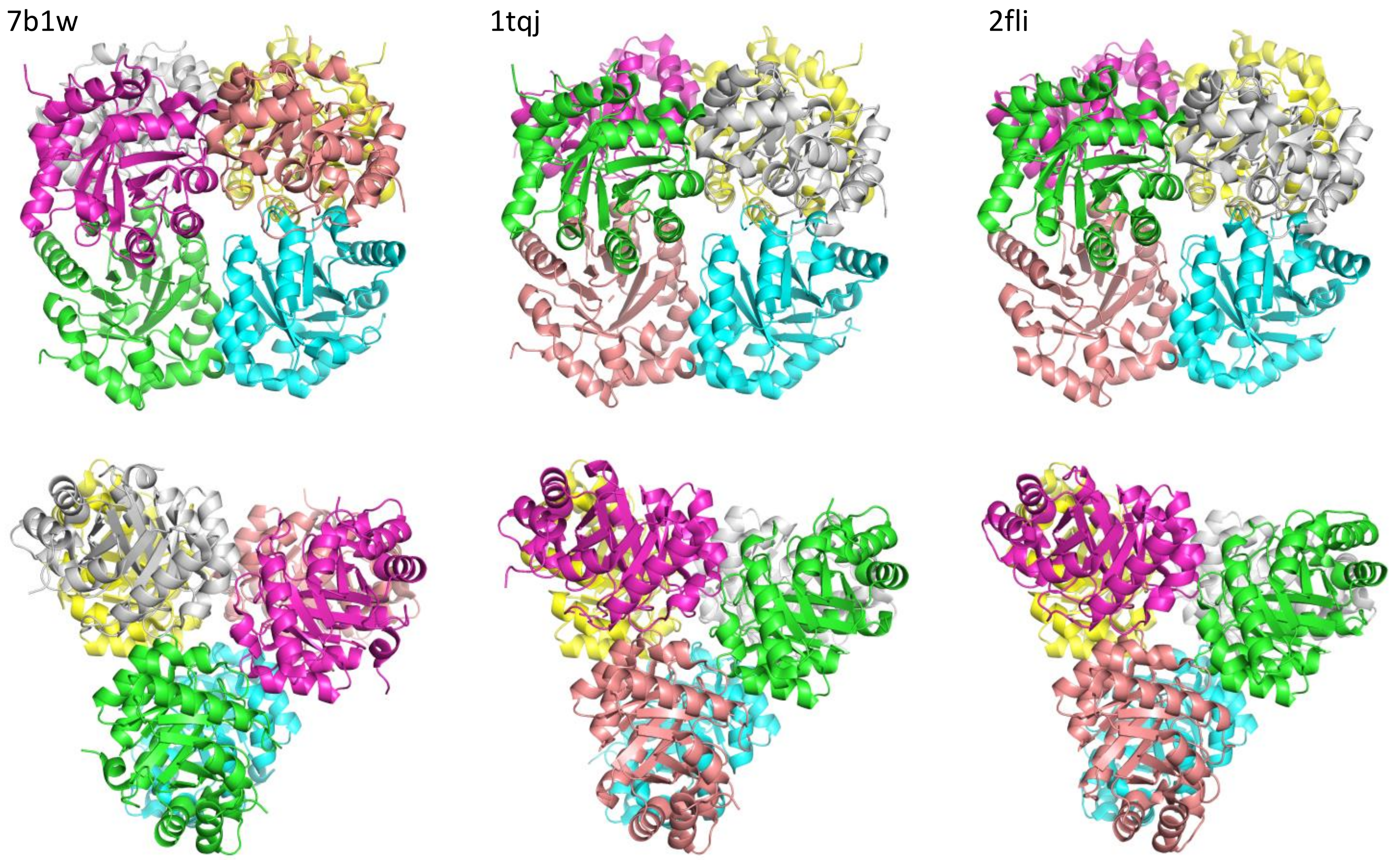

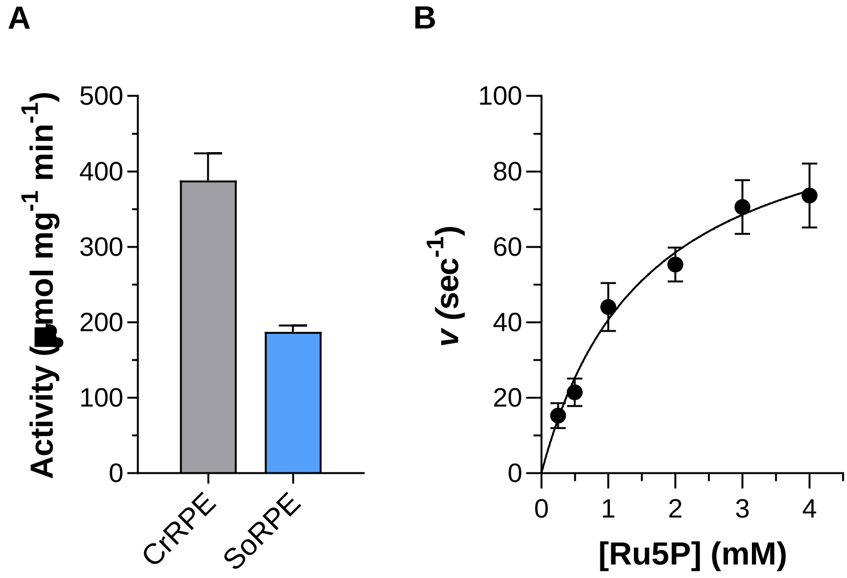
